# MiceVAPORDot: A novel automated approach for high-throughput behavioral characterization during E-cigarette exposure in mice

**DOI:** 10.1101/2023.10.27.564133

**Authors:** Yaning Han, Zhibin Xu, Zhizhun Mo, HongYi Huang, Zehong Wu, Xingtao Jiang, Ye Tian, Liping Wang, Pengfei Wei, Zuxin Chen, Xin-an Liu

## Abstract

The prosperity of electronic nicotine delivery systems (ENDS) or e-cigarette use has been regarded to lead an increasing risk of nicotine addiction, especially among youth. Understanding and evaluating the behaviors induced by ENDS are fundamental to the study of neuropsychiatric effects of e-cigarettes. However, little is known regarding the behavioral features during e-cigarette exposure in mice. Current behavioral assessments for nicotine addiction are based on nicotine withdrawal-induced anxiety which can be only performed after ENDS exposures. Here we developed MiceVAPORDot, a novel high-throughput tool for automated *in situ* behavioral characterization during e-cigarette exposure. The integration of a deep learning-based animal pose tracking method by MiceVAPORDot allows precise characterization on behavioral phenotypes of e-vapor exposed mice, which were unable revealed by traditional evaluation methodology such as conditioned place preference and elevated plus maze tests. The behavioral fingerprints recognized by MiceVAPORDot can be used for high-throughput screening on incentive nature of e-cigarette flavors as well as medications for smoking cessation.

## Introduction

Smoking is the leading cause of preventable death worldwide. Almost 7 million people died from tobacco use globally each year (Ahluwalia et al., 2018; Reitsma et al., 2017). Electronic nicotine delivery systems (ENDS) or e-cigarettes (e-cig) were first produced in 2003 as a safer alternative to combustible cigarette. E-cig delivers nicotine by vaporizing nicotine containing oil, which commonly combined with liquid flavors, instead of burning tobacco leaves. However, e-cig use or vaping, has dramatically increased in popularity, especially among adolescents, and potentially convert the new generation of non-smokers into smokers (Keller-Hamilton et al., 2021; Kinnunen et al., 2021; Tackett et al., 2021). Emerging researches start to focus on the identification of e-cigarette, or vaping product use-associated lung injury (EVALI) (Perrine et al., 2019). More than 2800 cases of hospitalized EVALI or deaths have been reported to the Centers for Disease Control and Prevention (CDC) in US by February, 2020 (Centers for Disease Control and Prevention. Outbreak of Lung Injury Associated with the use of e-cigarette, or vaping products. www.cdc.gov/EVALI. Accessed August 24, 2020.). Therefore, systematic and comprehensive approaches are needed for further investigation on the ENDS integrating epidemiologic, clinical, and laboratory data. The focus and scope of actions needed to address ENDS are grounded in not only the research illustrating the e-cig associated injuries and diseases but also the approaches to breaking e-cig addiction.

Nicotine is the major addictive component in both tobacco and e-cigs, which plays a critical role in the initiation and maintenance of smoking or vaping. Most of the knowledge about how nicotine affects behaviors in animals by using systemic injection regime of nicotine such as subcutaneous injection, osmotic pumps, or intraveneous self-administration. However, our understanding of the characteristic behavioral phenotypes induced by e-cig aerosol exposure in mice has remained rudimentary.

Nicotine addiction, similar to other substance use disorders, is a chronic disorder characterized by compulsive seeking behavior, loss of intake control, and the emergence of a “negative emotional state”. Behavioral assessments that have been widely used to evaluate the extent of nicotine addiction include daily water consumption measurement, conditioned place preference (CPP), as well as behavioral tests that evaluate nicotine withdrawal syndromes such as anxiety-like behaviors when nicotine is withdrawn (O’Dell et al., 2006; Winger et al., 2005). Dynamic progression from abuse to dependence which involves the changes of neurocircuits that affect rewarding and aversive systems in the brain is critical to understand the effects of nicotine in vapor or e-cig, and to characterizing mechanistic targets for countering nicotine dependence. Video-based analysis of ENDS addiction in animal models will undoubtedly drive future studies regarding comparions on addictive nature of e-cigarette flavors as well as medications for smoking/e-cig cessation.

Here, we developed the MiceVAPORDot, a novel behavioral assessment approach for e-cigarette addiction in mice based on *in situ* behavioral analysis during aerosol exposure inside the e-cig chamber. MiceVAPORDot, a deep-learning based data-driven analysis allows high-throughput pose tracking for each single mouse. Seven behavioral parameters were defined to quantify their location preference inside the chamber as well as the body postures, by which we may identify the different behavioral phenotypes between the mice exposed to e-cig aerosols without nicotine [E-cig(−Nic)] or with nicotine [E-cig(+Nic)]. Our current results demonstrated unique behavioral phenotypes in mice with different time length of nicotine-containing e-cig exposures. These behavioral changes were not able to be characterized by the traditional behavioral tests including conditional place preference and elevated plus maze. Further, the tensor component analysis (TCA) demixes the behavioral fingerprints of nicotine from the complex combination of behavioral parameters. The behavioral fingerprints to ENDS users in mouse models and facilitate rewarding/addictive assessments on e-cig products.

## Results

### MiceVAPORDot is an automated tool for high-throughput behavioral characterization in mice during e-cigarette nicotine exposure

The traditional evaluation strategies to quantify the behavior induced by addictive substances include behavioral evaluations in rodents including non-contingent models in which animals are passively exposed to rewarding substances, as well as widely used contingent models such as drug self-administration and relapse (Charbogne et al., 2014; X. Y. Hu et al., 2022; Lüscher et al., 2020; O’Dell & Khroyan, 2009), however the precise behavioral assessments during drug self-administration or taking is lacking, especially the behavioural profile while smoking. Here we developed MiceVAPORDot, which enabled the real-time monitor of the posture dynamics during aerosol exposures MiceVAPORDot(Fig. 1A). The main instrument of MiceVAPORDot is a sealed chamber with a aerosol delivery pump, which simulates the smoking condition of human. The pump delivers aerosol with precise concentration through a tube connected to the center of the chamber, and the aerosol is released from a circular outlet. Eight Dividers separates the chamber space equally into eight separate symmetrical flabellate areas. Mice were put into each flabellate area for simultaneous behavioral monitoring while “e-cigarette smoking”. The time-varying posture dynamics was achieved by a camera on the top of the chambers. The captured video frame without aerosol is demonstrated in Fig. 1B, and the frame with aerosol is demonstrated in Fig. 1C. Indeed, the vapor exposure program established in the inExpose e-cigarette device was to simulated human smoking habits and comply with the International Organization for Standardization (ISO) standard. Each puff consisted of a 30-second cycle, including 3 seconds for vaporization and 27 seconds of rest. The volume of each puff was set at 55 mL, and a bias flow of 2 L/min was maintained. The detailed configuration refers to Table S1. The urine level cotinine, the metabolite of nicotine, as measured in vapor exposed mice have confirmed the effective delivery of nicotine in our experiment (Fig. S1).

**Fig. 1.**
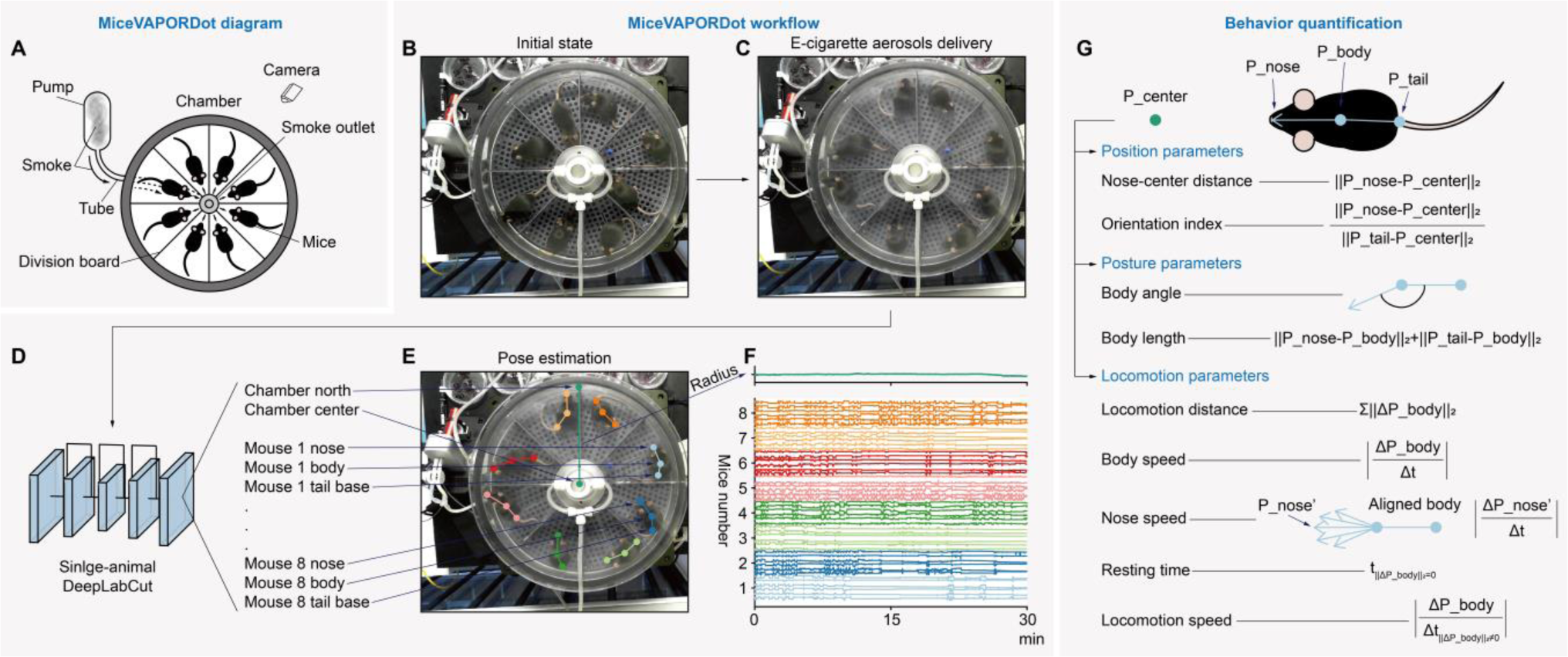
The framework of MiceVAPORDot. **A**, The hardware diagram of MiceVAPORDot. **B**, The initial state of MiceVAPORDot. Mice are put into MiceVAPORDot and ready for the smoke. **C**, The e-cigarette aerosol delivery of MiceVAPORDot. **D**, Using single-animal DeepLabCut to track key points of MiceVAPORDot. The chamber key points include the north and center points. The key points of each mouse include the nose, body, and tail base. **E**, The mice pose estimation and key points tracking with smoke. **F**, The tracking trajectories. Top, the radius is calculated to transform MiceVAPORDot from camera coordinates to world coordinates. Bottom, the coordinates of each mouse. The trajectories are ranked according to the sequence of the nose, body, and tail base. **G**, The behavior quantification of each mouse. The behavioral biomarkers are quantified into position, posture, and locomotion parameters. The position parameters include nose-center distance and orientation index. The posture parameters include body angle and body length. The locomotion parameters include locomotion distance, body speed, nose speed, resting time, and locomotion speed. (P_center, the center coordinate points of the chamber. P_nose, the coordinate points of the mouse nose. P_tail, the coordinate points of the mouse tail base. ||•||_2_, binormal operation. |•|, absolute value operation. Δ, differentiate operation. Σ, summation operation.).

The premise of behavior quantification of mice is pose-estimation, which means the definition and tracking the key nodes of the body of animals (M. W. Mathis & Mathis, 2020). In MiceVAPORDot, the heterogeneously diffused aerosol could cause complex occlusion of mice in video frames. In this case, most classical computer vision-based animal tracking approaches lack the capability to analyze the behavior of vapor exposed animals due to the complex environmental background (De Chaumont et al., 2012). Therefore, here we used an emerging toolbox called DeepLabCut (A. Mathis et al., 2018) to track the animals in MiceVAPORDot (Fig. 1D), which is a deep-learning based method for identification, tracking, pose estimation, and behavior classification of animals in complex environments (A. Mathis et al., 2018; Pereira et al., 2018).

First, MiceVAPORDotour data suggested that the single-animal pose estimation, instead of the multi-animal version of DeepLabCut was sufficient to accomplish the multi-animal tracking task in the MiceVAPORDot (Lauer et al., 2022). The active zone of all mice in the MiceVAPORDot are separated, therefore the location-scale distribution of each mouse could be accurately distinguished in DeepLabCut. Single-animal pose estimation model was able to process these mice as different animals although they look like each other. The model could learn separated distributions of mice and avoid identity errors. The advantage of using single-instead of multi-animal pose estimation is the high processing efficiency, which omits the steps of part affinity fields and identification (Lauer et al., 2022). The tracking results in our experiments are illustrated in Fig. 1E.

In order to calculate the physical size of mice, two points of the MiceVAPORDot were tracked including the chamber north (the north point of the MiceVAPORDot, Fig. 1D and E) and chamber center (the central point of the MiceVAPORDot, Fig. 1D and E), besides the real-time tracking of the nose, body, and tail base of each mouse. The physical radius of the chamber is 12 cm, and the pixel radius in each frame was calculated by the points of chamber north and chamber center. The transform scale from the camera to the real world was estimated by using the physical radius and pixel radius (Fig. 1F) and the trajectories of each mouse was identified and transformed from the video data to the actual coordinates.

Key-point trajectories are generated as high-dimensional time series, which are complex scenarios (Han et al., 2022). To establish a user-friendly database for behaviors of all the mice in vapor exposure, the trajectories are transformed into two groups of parameters to depict the location preferences and body postures of mice (Fig. 1G). In the MiceVAPORDot, the position parameters include the nose-center distance and the orientation index. The nose-center distance represents the desire of the mouse approaching the vapor outlet which can be used as a quantitative evaluation indicator of rewarding effects of the vapor. The orientation index is a relative value by comparing the nose-center distance and the tail base-center distance to quantify the degree of body orientation of mice toward the center of the chamber. Meanwhile, the posture parameters which were defined and applied in our experiment include body angle and body length (Taki et al., 2013). The locomotion parameters include locomotion distance, body speed, nose speed, resting time, and locomotion speed.

In order to demonstrate the advantage of MiceVAPORDot on behavioral monitoring of vapor exposed animals, we designed a long-term experimental paradigm that lasted for ∼50 days (Fig. 2). The mice were randomly divided into two groups including e-cigarette aerosols without nicotine [E-cig(−Nic)], and e-cigarette aerosols with nicotine [E-cig(+Nic)]. The mice were administrated with e-cigarette aerosol exposures in MiceVAPORDot for seven consecutive days (Fig. 2A), and followed by EPM test to assess their anxiety-like behaviors (Fig. 2B). After seven times of e-cigarette aerosol exposure in MiceVAPORDot, the mice received no exposure for the following 15 days (Fig. 2C). The CPP test was performed during the final three days of e-cigarette aerosol exposures along with behavioral observations in MiceVAPORDot.

**Fig. 2.**
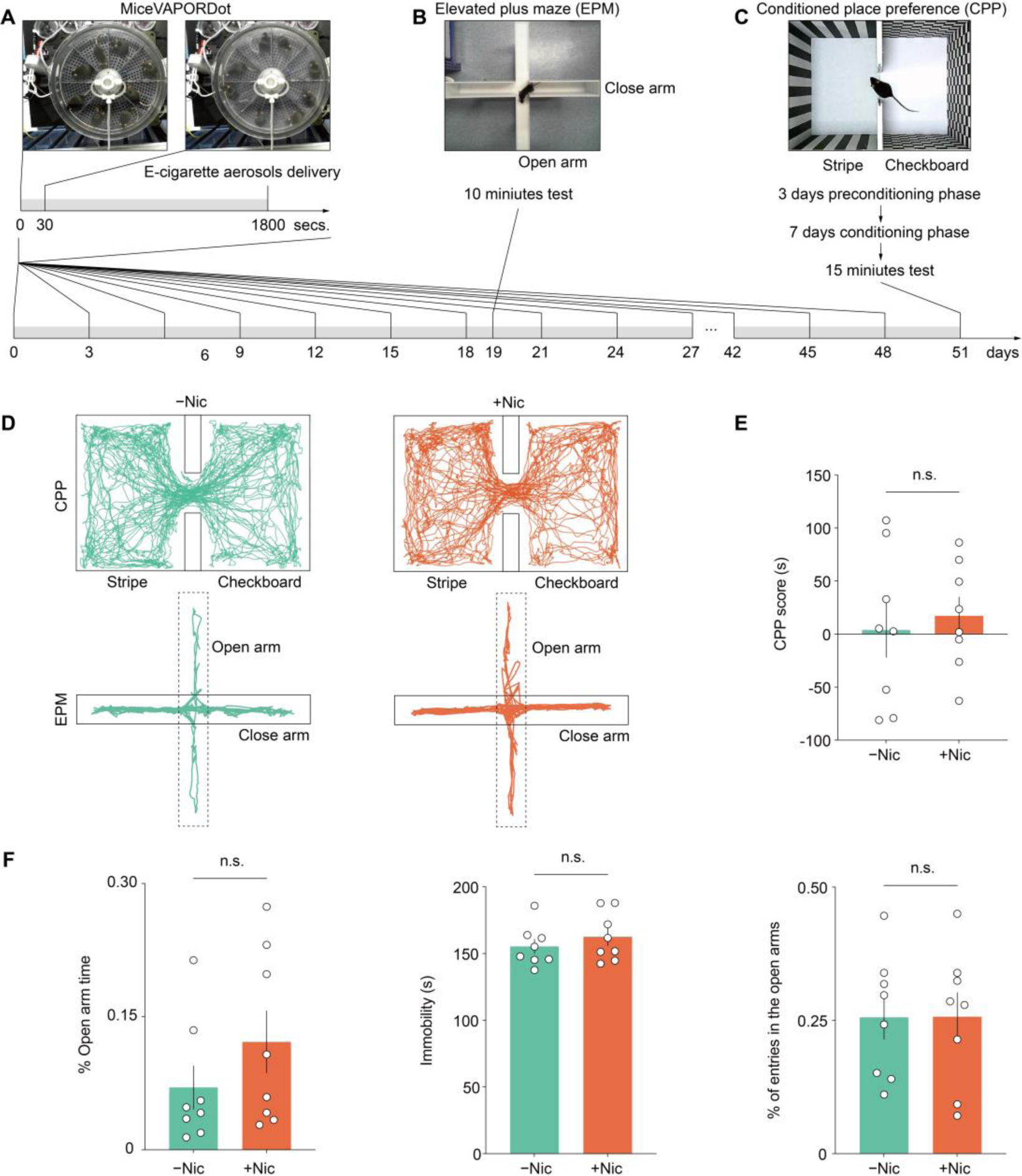
The experimental paradigms and the behavioral comparison of Conditioned Place Preference (CPP) and Elevated Plus Maze (EPM). **A**, The experimental paradigm of MiceVAPORDot. **B**, The experimental paradigm of elevated plus maze. **C**, The experimental paradigm of conditioned place preference. D, Trajectories of the -Nic (left) and +Nic (right) groups in the CPP test (upper) and EPM test (lower). E, No significant difference was found in the CPP score between the -Nic and +Nic groups. F, The results of the percent of open arm time, immobility time, and the ratio of Open/Total arm entries showed no significant difference.

### No behavioral phenotype after long term nicotine aerosol exposures was detected by traditional behavioral tests

Traditional behavioral tests such as CPP and EPM were generally used to assess the neuropsychiatric effects of nicotine. Nicotine’s rewarding effects are usually measured by CPP test and the EPM are performed to evaluate the anxiety-like behaivors during withdrawal period. In the CPP and EPM tests of the vapor exposed mice, the trajectories of mice were obtained from captured videos (Fig. 3A), from which the CPP preference score did not show significant differences between the two groups (Fig. 3B). Further, no statistically significant differences on the the ratio of mice entering the open arms and the closed arms in the EPM were observed between the e-cig vapor exposed groups with or without nicotine. Therefore no behavioral phenotype induced by nicotine aerosol exposures was detected by traditional behavioral tests such as the CPP and EPM tests under our nicotine vapor exposure conditions.

**Fig. 3.**
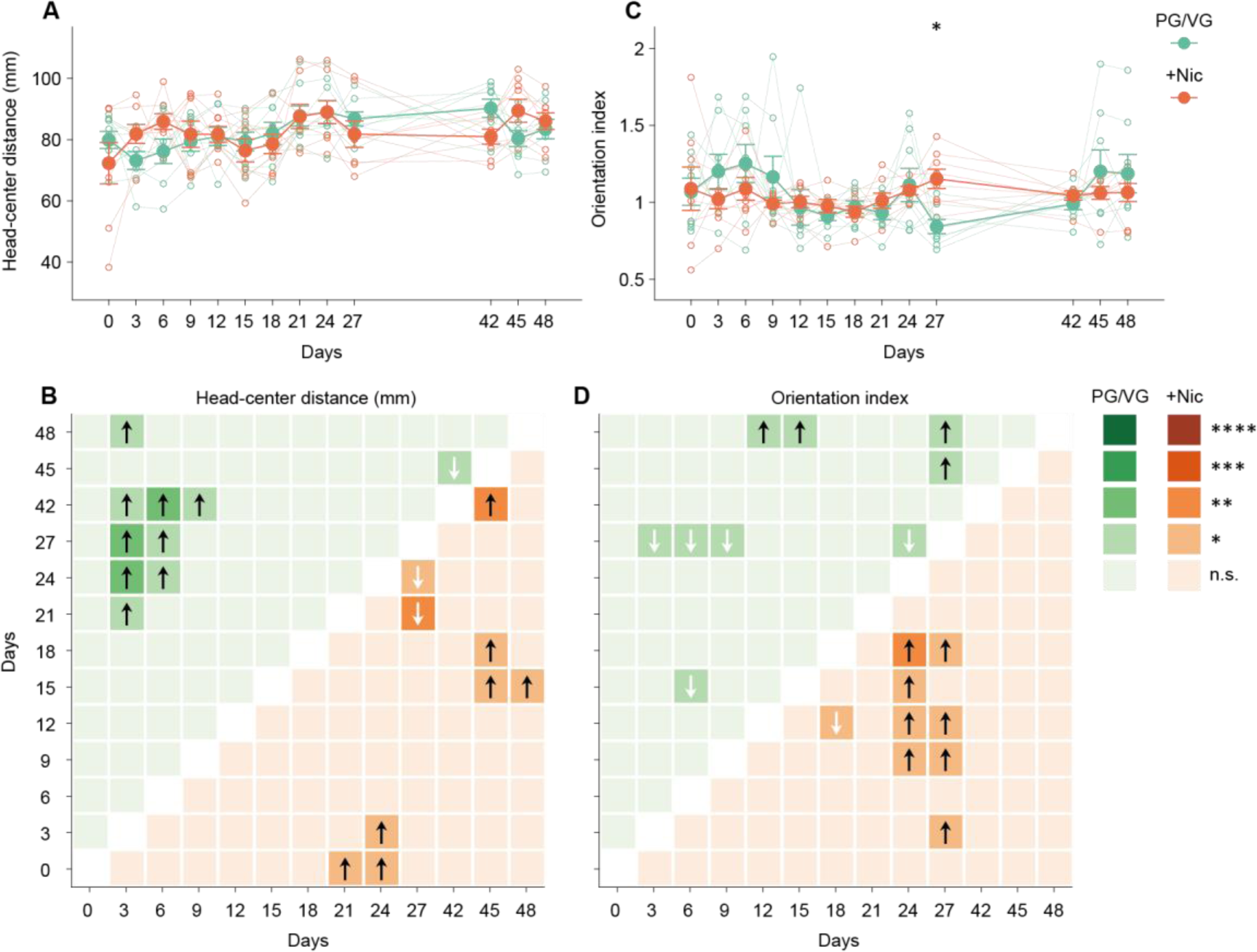
Alterations on position parameters in PG/VG and +Nic groups. **A**, The head-center distance shows no significant differences between two groups. **B**, The head-center distance of two groups shows opposite significant differences between 42 and 45 days. **C**, The orientation index of the +Nic group is significantly higher than PG/VG group in 27 day. **D**, The orientation index of two groups shows opposite significant differences between the 27 days. (****P < 0.0001. ***P < 0.001. ∗∗P < 0.01. ∗P < 0.05. n.s., no significant difference.).

### Mice show position preference after aerosol exposure in MiceVAPORDot

The position preference is described by head-center distance and orientation index. Comparing the position preference of the two groups, the +Nic group shows significantly increase of orientation index in the 27 day (Fig. 3A and B). Considering that the individual differences are large in nicotine exposure, we take a strategy to regard each animal as the baseline of themselves to compare the behavioral changing across days (Fig. 3B, 3D, 4B, 4D, 5B, 5D, 5F, 5H, and 5J). Here we call the comparison as ‘checkboard tiles’ for the convenient statement in later text. These comparisons indicate more subtle differences between two groups. The comparisons can be divided into the number of significant increase, the number of significant decrease, the number of total significant changes, the significant changes between different days.

**Fig. 4.**
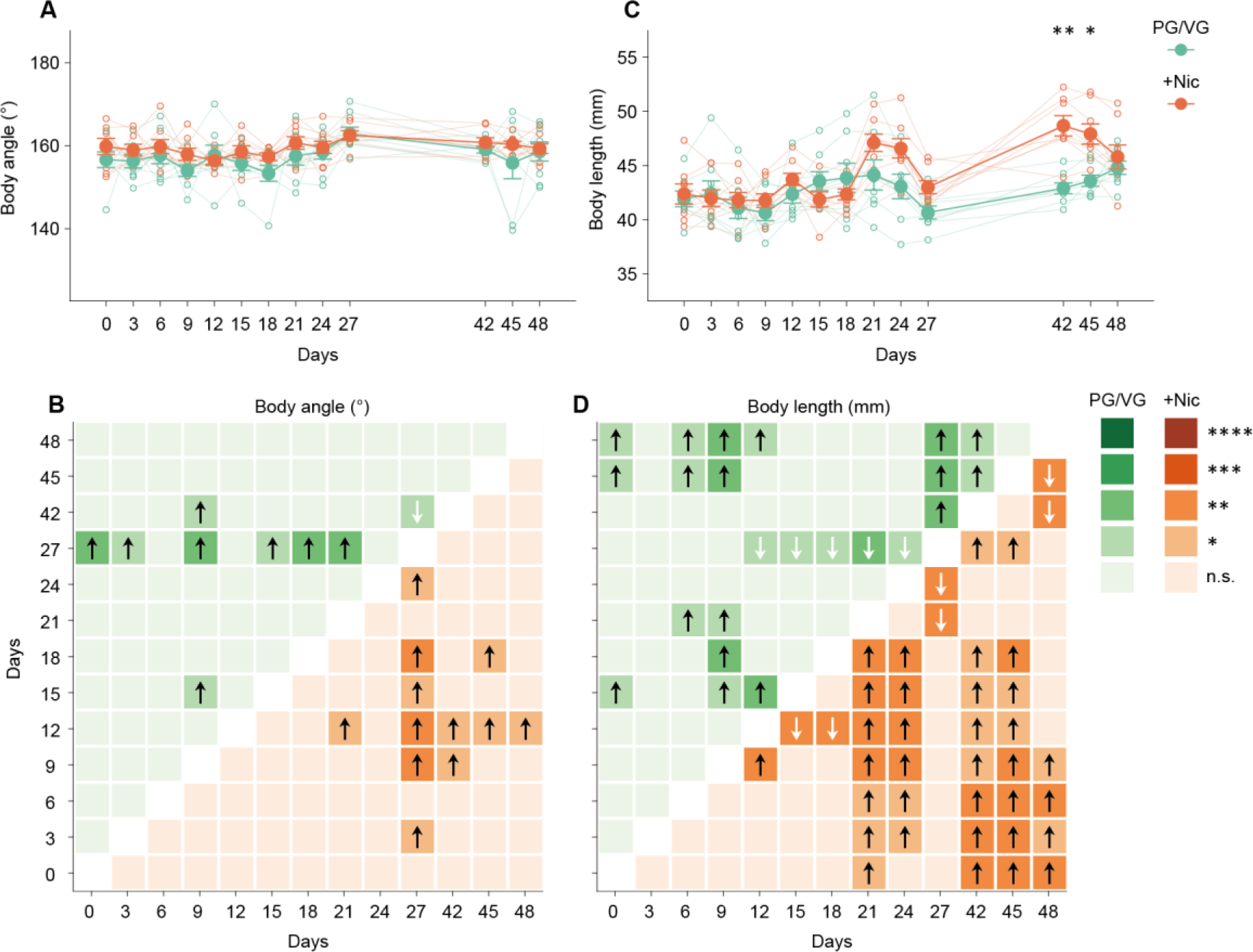
Alterations of posture parameters in PG/VG and +Nic groups. **A**, The body angle shows no significant differences between two groups. **B**, The body angle of the +Nic group shows more significantly increase in 42 to 48 days. **C**, The body length of the +Nic group is significantly higher than PG/VG group in 42 and 45 days. **D**, The body length of the +Nic group shows more significantly increase than the PG/VG group. (****P < 0.0001. ***P < 0.001. ∗∗P < 0.01. ∗P < 0.05. n.s., no significant difference.).

The number of significant increase of head-center distance of PG/VG group is 9 and +Nic group is 7 (Fig. 3B). The PG/VG group has 2 more significant increase of head-center distance than the +Nic group. The number of significant decrease of head-center distance is 1 versus 2. The PG/VG group has 1 less significant decrease of head-center distance than the +Nic group. The number of total significant changes is 8 increase versus 5 increase. The PG/VG group has 3 more total increase than +Nic group. The significant differences in 42-45D (42 compared with 45 days) are opposite between two groups. The PG/VG group shows significant decrease but the +Nic group shows significant increase. These comparisons illustrate more potential differences of head-center distance which cannot be revealed in Fig. 3A.

Although the orientation index only shows significant difference between two groups in the day 27 (Fig. 3C), the checkboard tiles display more subtle differences of these two groups (Fig. 3D). The number of significant increase of orientation index of PG/VG group is 4 and +Nic group is 8. The PG/VG group has 4 less significant increase of orientation index than the +Nic group. The number of significant decrease of orientation index is 5 versus 1. The PG/VG group has 4 more significant decrease of head-center distance than the +Nic group. The number of total significant changes is 1 decrease versus 7 increase. The PG/VG group has 8 less total increase than +Nic group. The increase of orientation index of +Nic group mostly in day 24 and 27, but the PG/VG group shows decrease in day 27.

In summary, these results show that the changing of position preference between PG/VG and +Nic groups are different. The changing of position preference occurs in about day 21, and the changing of position preference of the PG/VG and +Nic groups are opposite in day 21, 24, 27 and 45.

### The increase body length of mice after nicotine aerosol exposure

The posture of mice is described by body angle and body length parameters. Although the comparison of body angle shows no significant differences between two groups (Fig. 4A), the checkboard tiles show different changes of body angle of the two groups (Fig. 4B). The number of significant increase of body angle of PG/VG group is 8 and +Nic group is 12. The PG/VG group has 4 less significant increase of body angle than the +Nic group. The number of significant decrease of body angle is 1 versus 0. The PG/VG group has 1 more significant decrease of body angle than the +Nic group. The number of total significant changes is 7 increase versus 12 increase. The PG/VG group has 5 less total increase than +Nic group. The time latency of body angle increase of +Nic group (day 21 to 48) is longer than PG/VG group (more in day 27).

The body length of +Nic group is significantly higher than PG/VG group in day 42 and 45 (Fig. 4C), and the checkboard tiles show more differences (Fig. 4D). The number of significant increase of body length of PG/VG group is 18 and +Nic group is 34. The PG/VG group has 16 less significant increase of body length than the +Nic group. The number of significant decrease of body length is 5 versus 6. The PG/VG group has 1 less significant decrease of body length than the +Nic group. The number of total significant changes is 13 increase versus 28 increase. The PG/VG group has 15 less total increase than +Nic group. The +Nic group shows more increase of body length than the PG/VG group in day 21 to 24 and 42 to 48. Nevertheless the +Nic group shows more early body length decrease than the PG/VG group in day 15 to 18.

In summary, the results show that the body postures significantly changed in day 42, which reflects in the body length postures. The body angle and body length both increase after the nicotine aerosol exposure.

### Altered locomotion patterns of mice after nicotine aerosol exposure

The locomotion of mice is described by locomotion distance, body speed, nose speed, resting time, and locomotion speed parameters. The locomotion distance of +Nic group is significantly higher than the PG/VG group in day 27 (Fig. 5A). The number of significant increase of locomotion distance of PG/VG group is 8 and +Nic group is 10 (Fig. 5B). The PG/VG group has 2 less significant increase of locomotion distance than the +Nic group. The number of significant decrease of locomotion distance is 8 versus 1. The PG/VG group has 7 more significant decrease of locomotion distance than the +Nic group. The number of total significant changes is 0 increase versus 9 increase. The PG/VG group has 9 less total increase than +Nic group. PG/VG group shows earlier decrease of locomotion distance in day 6. PG/VG group also shows more decrease of locomotion distance than +Nic group in day 24 to 42. The body speed of +Nic group is significantly higher than the PG/VG group in day 27 (Fig. 5C). The patterns of significant differences in checkboard tiles of body speed (Fig. 5D) are the same as the locomotion distance (Fig. 5C) because body speed is calculated using locomotion distance subdivided the total time. Therefore the checkboard tiles patterns of body speed are not described here.

**Fig. 5.**
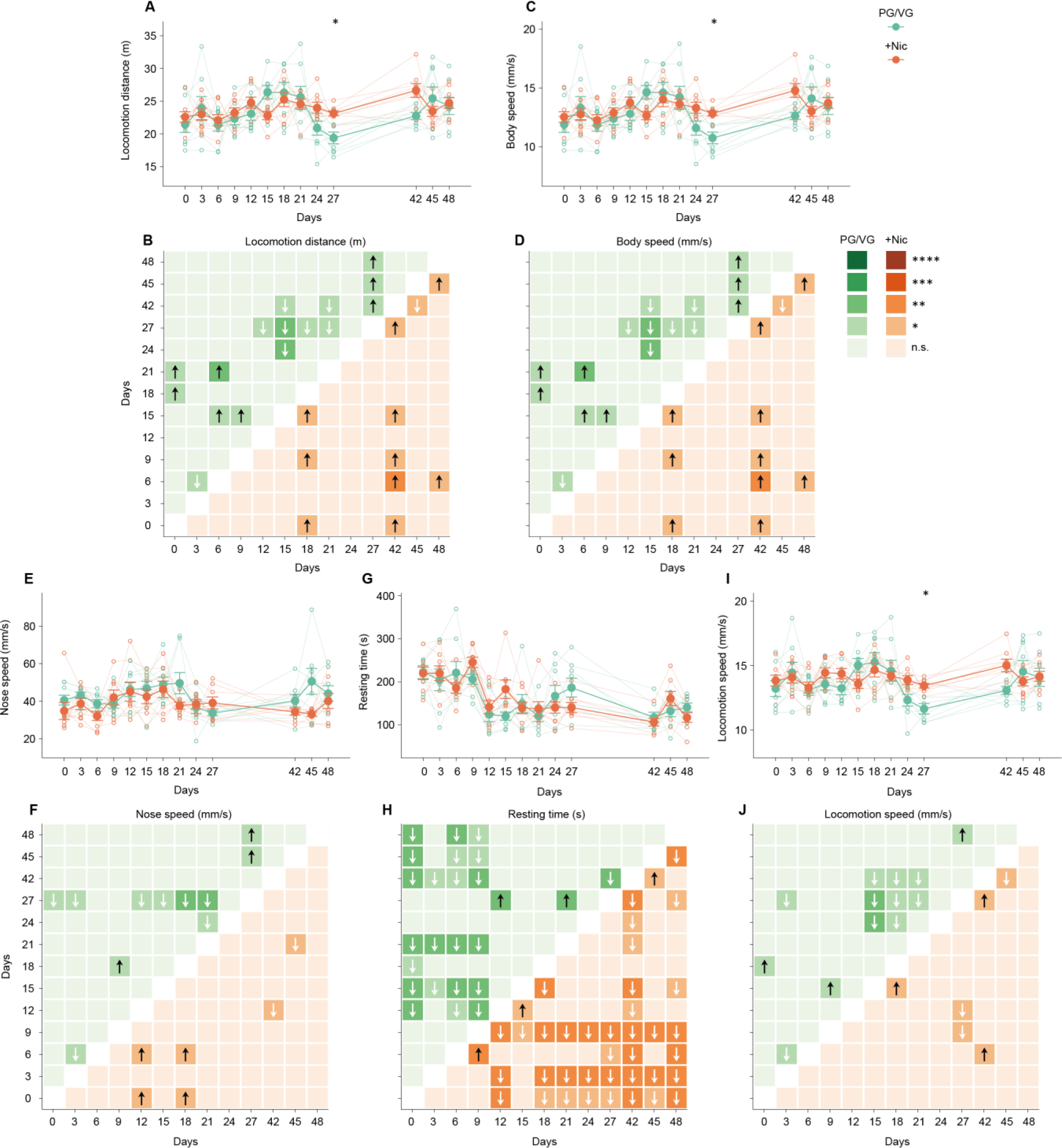
Alterations of locomotion parameters in PG/VG and +Nic groups. **A**, The locomotion distance of the +Nic group is significantly higher than PG/VG group in the 27 day. **B**, The locomotion distance of the PG/VG group shows more significantly decrease in the 24 to 42 days. **C**, The body speed of the +Nic group is significantly higher than PG/VG group in the 27 day. **D**, The body speed of the PG/VG group shows more significantly decrease in the 24 to 42 days. **E**, The nose speed shows no significant differences between two groups. **F**, The nose speed of the PG/VG group shows more significantly decrease in the 27 day. **G**, The resting time shows no significant differences between two groups. **H**, The resting time of the +Nic group shows more significantly decrease than the PG/VG group. **I**, The locomotion speed of the +Nic group is significantly higher than PG/VG group in the 27 day. **J**, The locomotion speed of the PG/VG group shows more significantly decrease than the +Nic group. (****P < 0.0001. ***P < 0.001. ∗∗P < 0.01. ∗P < 0.05. n.s., no significant difference.).

The nose speed of two groups shows no significant difference (Fig. 5E). The number of significant increase of nose speed of PG/VG group is 3 and +Nic group is 4 (Fig. 5F). The PG/VG group has 1 less significant increase of nose speed than the +Nic group. The number of significant decrease of nose speed is 8 versus 2. The PG/VG group has 6 more significant decrease of nose speed than the +Nic group. The number of total significant changes is 5 decrease versus 2 increase. The PG/VG group has 7 less total increase than +Nic group. The PG/VG group shows earlier decrease of nose speed than +Nic group in day 6. The +Nic group shows earlier increase of nose speed than +Nic group in day 12. The PG/VG group shows more significant decrease of nose speed than +Nic group in day 27, and more significant increase in day 45 to 48.

The resting time of two groups shows no significant difference (Fig. 5G). The number of significant increase of resting time of PG/VG group is 2 and +Nic group is 3 (Fig. 5H). The PG/VG group has 1 less significant increase of resting time than the +Nic group. The number of significant decrease of resting time is 23 versus 37. The PG/VG group has 14 more significant decrease of resting time than the +Nic group. The number of total significant changes is 21 decrease versus 34 decrease. The PG/VG group has 13 less total decrease than +Nic group. +Nic group shows earlier increase of resting time than the PG/VG group in day 9. In day 18 to 48, the +Nic group shows continuous decrease of resting time but the PG/VG group is interrupted in day 24 and 27.

The locomotion speed of +Nic group is significantly higher than the PG/VG group in day 27 (Fig. 5I). The number of significant increase of locomotion speed of PG/VG group is 3 and +Nic group is 3 (Fig. 5J). The locomotion speed increase of PG/VG group are the same as the +Nic group. The number of significant decrease of locomotion speed is 10 versus 3. The PG/VG group has 7 more significant decrease of locomotion speed than the +Nic group. The number of total significant changes is 7 decrease versus 0 decrease. The PG/VG group has 7 more total increase than +Nic group. The PG/VG group shows earlier decrease than the +Nic group in day 6. The PG/VG group shows more consistent decrease of locomotion speed than the +Nic group in day 24 to 42.

In summary, the locomotion of mice in +Nic group shows more increase than the PG/VG group. The locomotion distance, body speed, nose speed, and locomotion speed decrease in PG/VG group but increase in +Nic group, and the resting time increase in PG/VG group but decrease in +Nic group.

### The behavioral fingerprints for mice exposed to nicotine aerosol

The spontaneous locomotor activity and delicate behavioural characteristics recognized by AI based automated assessments have become a useful tool for behavioral fingerprint identification and drug screening (Datta, Nature Neuroscience 2020; Datta, Neuron, 2023). By using MiceVAPORDot, we have extracted a total of nine behavioral parameters in mice exposed to nicotine containing vapor, and these behavioral parameters are temporally dynamic. Here we have performed tensor component analysis (TCA) to extract the most significant responses to nicotine vapor among the complex combination of behavioral parameters (Williams, Neuron, 2018). First, we converted the raw data into a tensor that includes the information of individual mouse and the date (Fig. 6A), the non-negative constraints method was used to calculate the variability explanatory degree and reconstruction error according to the number of different components, so as to select the appropriate number of components for subsequent analyses (Fig. 6B-C). Our calculation results suggested that, 30 selected components were able to explain more than 90% of the variability, meanwhile, the reconstruction error was also significantly decreased and close to convergence at 30 components, therefore we selected the factor number of 30 for the subsequent analyses.

**Fig. 6.**
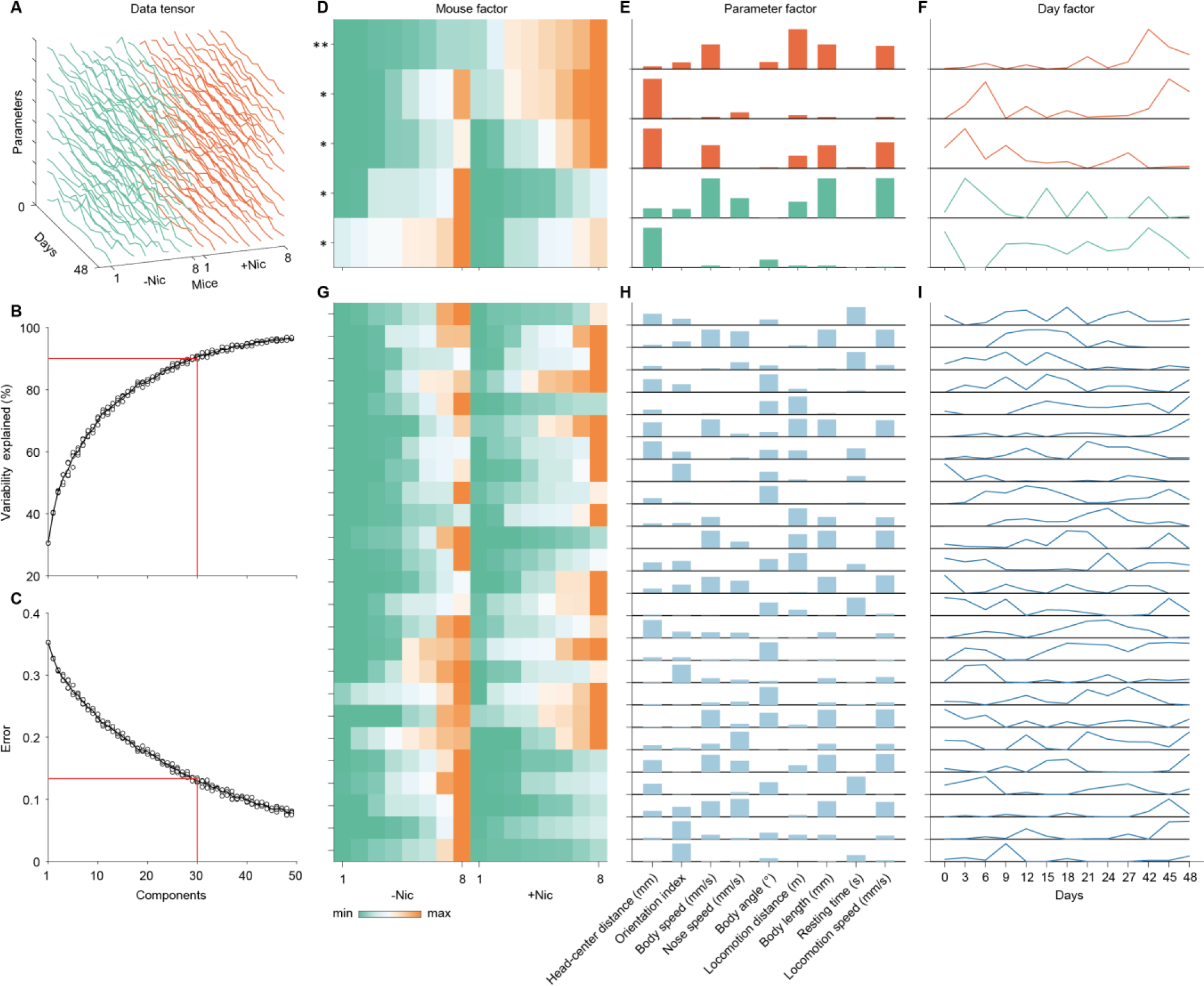
Tensor component analysis (TCA) demixes the behavioral fingerprints of nicotine exposure. **A**, All of the behavioral data are organized into a tensor. **B**, The percent variability explained by components during the optimization process of TCA. 30 components can explain more than 90% variability. **C**, The reconstruction error is not improve with the increase number of components. **D**, The five significantly different components of mouse factor between two groups. The components are sorted to illustrate the differences. **E**, The corresponding components of parameter factor of **D**. Orange bar represent the mean value of mouse factor components of +Nic group is larger than -Nic group. Green bar represents the mean value of mouse factor components of -Nic group is larger than +Nic group. **F**, The corresponding components of day factor of **D** and **E**. **G**, Other 25 mouse factor components with no significant differences. **H**, The corresponding parameter factor components of **G**. **I**, The corresponding day factor components of **G** and **H**. (∗P < 0.05. ∗∗P < 0.01.).

We then analysed the differences on the mouse factor caused by nicotine vapor exposure. We have calculated the differences in the components of mouse factor respectively, and found 5 components that have shown significant differences between PG/VG and +Nic groups (Fig. 6D), among which three of them have shown increased while the other two presented decreased levels in mice with +Nic group compared to PG/VG group. The most significant component of mouse factor can be regarded as the first effect of nicotine exposure. The PG/VG and +Nic group both show consistent trends in the component of mouse factor. The corresponding parameter factor can be separated into three groups according to the height of bars (Fig. 6E), which is the highest locomotion distance, and the medium group of body speed, body length, and locomotion speed, and the shortest other parameters. Combing with the day factor of this component, the most significant effect of nicotine exposure is the temporal increase of locomotion distance, which reaches the maximum value in day 42. The body speed, body length, and locomotion speed also show increase across time, although the increases takes less weights than locomotion distance. Other parameters do not change much in this component. The second significant component of mouse factor shows 1 outlier mouse in PG/VG group and 2 in +Nic group. The corresponding parameter factor only shows that the head-center distance takes the highest weight. The head-center distance shows two peaks in the day factor, respectively day 0 to 9 and day 42 to 48. The third significant component of mouse factor shows 1 outlier mouse in PG/VG group and 4 in +Nic group. The corresponding parameter factor also can be separated into three groups, which is the highest head-center distance, the medium body speed, locomotion distance, body length, and locomotion speed, and the other no change parameters. The day factor shows one high peak in day 0 to 6, and the trend of the day factor is decreasing. Some of the parameters of this component are overlapped with the first component, but the trends of their day factors are different. The reason is that this component is influenced by the individual difference. Some of the mice tend to show the decrease of these parameters of the third component, which is opposite to the main effect. The fourth significant component shows 1 highest outlier mouse in PG/VG group and the trends of +Nic group are more consistent. The corresponding parameter factor can be separated into three groups, respectively the highest body speed, body length, and locomotion speed group, the head-center distance, orientation index, nose speed, and locomotion distance group, and the other no change parameters as the third group. The day factor shows fluctuation with 4 peaks, in which the first peak is the highest. This component indicates that the changes of behavioral parameters of the PG/VG group are not stable. The fifth significant component of the mouse factor shows more similarity in the two groups. The head-center distance of the corresponding parameter factor is obviously higher than other parameters. The day factor shows the head-center distance in day 3 and 6 takes lowest weights. Additionally, the day factor of head-center distance decreases from day 0 to 6 and day 42 to 48. These results indicate that PG/VG-induced changing of position preference decreases in continuous exposure, and the position preference can be recovered after interval exposure.

The other 25 components of mouse factor with non-significant differences can explain the individual differences between two groups (Fig. 6G). The mouse factor of the 25 components indicates that there are about two mice show large individual differences inner each group. The parameter factor is not consistent in each component (Fig. 6H). Although most of the parameter factors (∼8 in 25) show only one distinct parameter such as head-center distance, orientation index, and locomotion distance in each component, the combination of parameters such as body speed, body length, and locomotion speed also exist. The day factor shows complex dynamics of each component (Fig. 6I). But in the global level, the day factor shows three peaks in day 0, day 15∼27, and day 42-28. They support the results of Fig 3, 4, and 5, which illustrate acute, long-term, and inter phases of the nicotine effects.

In summary, TCA demixes the combination of behavioral parameters in two groups. These behavioral combinations can be regarded as the fingerprints to indicate the nicotine exposure. Additionally, the behavioral fingerprints reveal the individual differences of each mice in each group, which can be a characteristic of each mouse during nicotine or PG/VG exposure.

## Discussion

MiceVAPORDot has the potential to be a powerful tool to fill the blank of assessing the behavior during the aerosol exposure procedure. Although previous studies have demonstrated that e-cigarette nicotine exposure could influence the behaviors of mice (Garrett et al., 2021; Grant et al., 2016; Ponzoni et al., 2020) no work have characterized behavioral alterations during smoke exposure. The key obstacle could be animal tracking under the occlusion of e-cigarette aerosols. Combing with the deep learning-based pose estimation method (A. Mathis et al., 2018), MiceVAPORDot enabled us to record dynamic behaviors of mice exposed to aerosol or smoke, and evaluated the subtle behavioral phenotypes induced by smoke exposure, which is the foundation of further analysis of neurobehavioral effects of atomized substances, such as nicotine or flavor containing e-cigarettes.

The distribution of data shows large individual differences in mice. A large number of previous studies have shown individual differences in sensitivity to nicotine (Hiroi & Scott, 2009; M. C. Hu et al., 2006). The rewarding effect of nicotine was reported species-conserved (Chellian et al., 2021; Palmer et al., 2021), we here used MiceVAPORDot to simulate human smoking conditions, and obtain the behavioral phenotypes “during smoking”. The behavioral analysis enabled us to dissect the behavioral characteristics corresponding to individual variability of nicotine sensitivity, facilitating further research on genetic vulnerability and gene-environment interactions in nicotine dependence, such as sex, history of other drug use or psychiatric disorders, and parental smoking.

Here we demonstrated that MiceVAPORDot was able to capture nicotine-related behavioral changes in mice that could not be detected by traditional behavioral tests. One of the reasons could be that significant behavioral phenotypes induced by nicotine are most obvious during the early stage of e-cigarette aerosol exposure but not later. This phenomenon is also common in humans. It is hard to distinguish nicotine-dependent persons according to their daily behaviors without smoke. Another reason could be that individuals have variability in initial nicotine sensitivity. Human studies have shown that only about half of daily smokers meet the criteria for nicotine dependence (M. C. Hu et al., 2006). It could be an explanation of the trend of the EPM preference ratio while no significant phenotypes were captured by traditional behavioral assessments in the case that the sample size was small in our experiment.

Here, by using MiceVAPORDot, we have observed the previously unrevealed position preferences induced by nicotine. Our data suggested that mice in both two groups showed escape-avoidance behaviors to the aerosol which was indicated by the increase of head-center distance. However, after the repeated aerosol exposures (about 27 days), the mice in PG/VG group tended to stay farther away from the aerosol outlet than the mice of +Nic group. These results demonstrate that prolonged nicotine-containing aerosol exposure reduced aversion to the aerosol in mice. It could also be further verified by the observations in the intermittent exposures of day 27 and 42. The head-center distance of mice on day 42 were longer than that on day 27 in PG/VG group but not in the +Nic group. Therefore, if only consider the changing of position preferences of +Nic group, the dislike of aerosol of mice would cover the phenomenon of the addictive effect of nicotine. Another behavioral phenomenon induced by nicotine aerosol captured by MiceVAPORDot was the significant increase of orientation index in mice of +Nic group, which can be explained by the differences in the concentration of aerosols in the chamber. The diffusion speed of smoke would continuously decrease from the smoke outlet to the edge of the chamber. It causes the higher concentration of aerosol at the edge of the chamber. The mice in +Nic group tend to face to the edge of the chamber to potentially take more nicotine from the location spot with higher concentration aerosol when observed from the raw videos. These results indicated that precise collection of behavioral parameters can provide a comprehensive understanding of neuropsychiatric effects of nicotine aerosol, which is the advantage of MiceVAPORDot.

The increase of body length was the main phenotype in body postures of mice during nicotine aerosol exposure was. Previous studies have shown that nicotine and smoke impaired the development of bone and reduced the body height (Durham et al., 2019; Ravnborg et al., 2011). Therefore, the increase of body length of mice in +Nic group in our experiments could not be attributed to its effect on development. Another explanation of the increased body length was that nicotine directly increases GABAergic transmission mediated by nAChRs on the GABA neurons (Laviolette & Van Der Kooy, 2004; Mansvelder et al., 2002) The induced action potential firing in GABA neurons allow the mice to enter a relaxed state (Lindstrom, 1997; Qin et al., 2022; Rupprecht et al., 2022), which was manifested by the increased body length. The increase of body angle in mice in +Nic group was another evidence indicating a relaxed state of mice exposed to nicotine aerosol. This phenomenon is species-conserved and can also be observed in smokers(Benowitz, 2008). Humans show relaxing postures during smoking, which is similar to the mice exposed to nicotine aerosol. Interestingly, the relaxing postures could be observed during nicotine aerosol exposure which was rarely presented after aerosol exposure in mice. Besides, the increase of body length mainly presented in day 21 and during aerosol exposures after day 42, confirming the relaxing effects induced by long-term aerosol exposure and re-exposure to nicotine. Similar phenotype can be observed in human smokers, in which the aversive behaviors were presented in first cigarette use and get addicted to nicotine after repeated episodes of nicotine exposure. The reward sensitization by nicotine is achieved after the withdrawal period. These results have convince the application value of MiceVAPORDot, that enables the dynamic dissections on neuropsychiatric behavioral alterations induced by e-cigarette aerosol, and can be used as a tool for screening for smoking cessation medications which are promising to be translated to humans.

Consistent with previous reports on effects of intravenous (Lenoir et al., 2013) and stereotactic injection (Panagis et al., 1996) of nicotine, another interesting behavioral alteration in nicotine aerosol exposed mice is the enhanced locomotion. This effect can be explained by increased activities of midbrain dopamine neurons induced by nicotine (Dongelmans et al., 2021; Fan et al., 2023). In sum, instead of quantifying one behavioral state by one behavioral paradigm in mice, the multiple behavioral modules captured by MiceVAPORDot enabled us to depict the comprehensive concomitant behavioral characteristics during aerosol exposure (Grunberg et al., 1988; Veltman et al., 1998). Further, we have utilized the TCA to reveal the potential weights of combinations of all the behavioral parameters. The higher weights of concomitant posture and locomotion parameters further revealed the simultaneous effects of nicotine as relaxant and excitant to mice during aerosol exposure. TCA in our study also explained the individual behavioral differences in mice caused by nicotine. The effects of nicotine are complicated, and related behavior changing could also be complicated. Considering the diversity of nicotine receptors in the body and the complexity of nicotine’s physiological effects, more precise behavioral characteristics urge to be uncovered by utilizing advanced analytical methods to decompose complex interactions between different behavioral variables. These demixed variables can be used as the behavioral fingerprints to build up a database for the comprehensive comparison in various experimental settings.

As a useful behavioral screening tool for the characterization of neurobehavioral effects of atomized substances, MiceVAPORDot can be further improved in three aspects. The first aspect is the control of the diffusion of smoke. Each area of MiceVAPORDot is flabellate, which makes the diffusion of smoke become unpredictable. In this case, it is hard to estimate the interactions of aerosol with mice in the chamber. The rectangle area can be a more reasonable alternative solution. Secondly, neural signal recording from certain neuronal populations in specific brain regions of mice during vapor exposure will further facilitate our understanding on the real-time neuropsychiatric actions of atomized substances corresponding to the behavioral fingerprints.

Since the current version of MiceVAPORDot could not allow the mice to implant neural recording devices which was due to the restriction by the chamber space, increasing the height of the division board of the Inexpose chamber will be our next step so as to add the wireless devices for neural recording and modulation (Cai et al., 2022). Finally, we will further improve the purely data-driven analysis on characteristic behavioral phenotypes of animals during aerosol/smoke exposure by utilizing the unsupervised behavior decomposition methods. Finer behaviors may reflect more subtle behavioral phenotypes induced by the treatments of nebulized drugs (Pereira et al., 2020). More cameras in different directions can be added to MiceVAPORDot for the 3D reconstruction of each mouse (Han et al., 2022). Additionally, the behavioral phenotype profile can be a more comprehensive description of the effects of e-cigarettes (Huang et al., 2021; Liu et al., 2021), the accurate tracking of animals requires massive dataset annotations for training the deep learning model which can be simplified by generative data annotation methods to increase the speed of data processing (Han et al., 2023).

In summary, although various novel model in mice with relevance to e-cigarette use bear great potential for the validation of potential addictive properties and harmful effects of nicotine vapor administration, our present work have established a novel tool for real-time behavioral analysis of mouse models during different periods of smoke exposure. The characteristic behavioral phenotypes will enable us reveal behavioral characteristics associated with e-cigarette addiction that have never been appreciated before. MiceVAPORDot developed in this work can facilitate assessments on atomized substances such as the neuropsychiatric effects of e-cigarettes with various flavors, it is also a valuable tool that can be extended to the unbiased evaluation on environmentally harmful substances. Our unbiased analyses through MiceVAPORDot may thereby advance the translational research field toward assessment approaches of drug nebulization therapies.

## Materials and Methods

### Animals

Male C57BL/6J mice (Hunan SJA laboratory animal Co., LTD. Hunan, China) aged 8 weeks old) were maintained in standard housing conditions on a 12/12 h day/night cycle with ad libitum access to food and water. All animal experiments and procedures respected the animal welfare guidelines to minimize the number and suffering of mice used and were approved by the Animal Care and Use Ethics Committee of the Shenzhen Institutes of Advanced Technology, Chinese Academy of Sciences (SIAT-IACUC-200828-NS-MZZ-A1431). All behavioral tests were conducted at a fixed period during the light cycle. All animal experiments were approved by the Animal Care and Use Committees at the Shenzhen Institute of Advanced Technology, Chinese Academy of Sciences.

### Apparatus and protocol of aerosol exposure

The MiceVAPORDot was developed based on the inExpose e-cigarette device (SCIREQ Scientific Respiratory Equipment Inc). The inExpose system, with its compact size and high level of integration, operates under a standard fume hood and offers the flexibility of both nose-only and/or whole-body exposure of rodents. It also features automated cigarette smoke and aerosol generation, making it a comprehensive and versatile research tool. A camera was strategically placed directly above the inExpose e-cigarette unit to allow video recording during experiments.

A commercially sourced E-cigarette liquid was used that consisted of 50% propylene glycol and 50% Glycerol (PG/VG) with or without nicotine at a 4% mass percentage concentration. As for delivery of e-cigarette aerosols, animals were randomly divided into 2 groups (n = 8) and exposed to the following conditions for around one month (Fig. 2A) during the model of addiction and behavior tests: e-cigarette aerosols without nicotine [E-cig(−Nic)], and e-cigarette aerosols with nicotine [Ecig(+Nic)]. Every single mouse in each group was placed in an independent chamber filled with e-cigarette aerosols once daily for 30 min as shown in Fig. 1A and Fig. 2A. The amount of nicotine of the aerosol in this study is equivalent to 30-min cigarette smoking daily based on nicotine content detection.

### Behavioral assessments

The behavioral assessments include elevated plus maze (EPM) (Rodgers & Dalvi, 1997) and conditional place preference (CPP) tests (Fudala et al., 1985).

To measure the anxiety levels of mice, the EPM was assessed by using a plastic elevated plus maze constructed from two white open arms (25 cm length × 5 cm width) and two white enclosed arms (25 cm length × 5 cm width × 15 cm height) extending from a central platform (5 cm length × 5 cm width) at 90° which form a plus shape (Fig. 2B). The maze was placed 65 cm above the floor. A camera was set directly above the EPM apparatus for video recording. The mice were individually placed at the center, with their heads facing the open arms. Moving trajectory and the amount of time spent in the same type of arms were recorded during the 10-min sessions and analyzed using customized MATLAB code.

An unbiased CPP test paradigm was used in this study (Fig. 2C). In brief, the place conditioning chambers consisted of two distinct compartments (40 cm length × 60 cm width × 40 cm height) with different geometric patterns, separated by a cross baffle that allowed access to either side of the chamber. In the preconditioning phase (before aerosols exposure), animals were allowed to move freely between the compartments for 15 min once daily for 3 days. Moving trajectory and time spent in each compartment was recorded as baseline measurements. These data were used to separate the animals into groups of approximately equal bias. The last 7 days in the model establish phase were the conditioning phase, during which all groups received e-cigarette aerosol exposure respectively and then were immediately confined to the same side of the compartment for 30 min. 4 hours later animals were confined in the opposite compartment for 30 min. After the last exposure, animals were immediately placed in a CPP compartment without cross baffle for 15 min. Moving trajectory and time spent in each compartment were analyzed using the Visualtrack software.

### Computational hardware and software

The data are processed by a workstation with one Intel i9-12900K CPU, one NVIDIA GeForce RTX 3090 GPU, 128 Giga Byte RAM, and 16 Tera Byte HDD. The deep learning animal pose estimation used DeepLabCut 2.2.1.1 version running on Python 3.8.13, which was installed using Anaconda. The analysis of behavioral biomarkers, CPP, and EPM used customized code running in MATLAB 2022b.

### Key-point tracking

The DeepLabCut was used for tracking all the key-points of mice (A. Mathis et al., 2018). In MiceVAPORDot, we defined a total of 26 key points including chamber north, chamber center, and nose, body, and tail base of each mouse (Fig. 1D). Totally 520 frames were manually labeled to train the deep learning-based pose estimation model ResNet50 (He et al., 2016). The iteration steps were set to 1030000, and other training parameters of the model were set to default. The final training loss is 0.00139, which is small enough to get high tracking performance (Huang et al., 2021). This well-trained model was used to track all the defined key points of the videos in this paper. The tracking results are appended to raw videos for the human check of tracking precision.

### Statistical analyses

All of the statistics were achieved by Prism 8.0 (GraphPad Software). Before hypothesis testing, data were first tested for normality by the Shapiro–Wilk normality test and for homoscedasticity by the F test. For normally distributed data with homogeneous variances, parametric tests were used (two-column data using unpaired two-tailed t-test, and two-column temporal paired data using paired two-tailed t-test); otherwise, non-parametric tests were used (two-column data using two-tailed Mann-Whitney test, and two column temporal paired data using two-tailed Wilcoxon test). The comparisons of data with one sample used the Freidman test with Dunn’s multiple comparisons test because the data illustrated in Fig. 5F and G do not pass the Shapiro–Wilk normality test.

## Acknowledgments

(optional; includes special thanks or dedications)

## Author contributions

(mandatory for each of your authors)

Zuxin Chen and Xin-an Liu designed the study; Yaning Han performed the behavioral analyses and wrote the original draft; Zhibin Xu conducted the majorities of aerosol exposure experiments and wrote the original draft; Zhizhun Mo, Zehong Wu and Xingtao Jiang participated in the aerosol exposure experiments. HongYi Huang and Ye Tian participated in the behavioral assessments. Liping Wang, Pengfei Wei, Zuxin Chen and Xin-an Liu revised the manuscript. All authors contributed to and have approved the manuscript.

## Funding

(This must cover all authors and sources of funding listed for the manuscript.)

This work was supported by grants from the STI2030-Major Projects (2022ZD0207100 to Z.X.C), the National Natural Science Foundation of China (NSFC) (32371213, 31900728 to X.A.L, 32000710 to Z.X.C, 32222036 to P.F.W, U20A2016 to Z.X.C), the Guangdong Basic and Applied Basic Research Foundation (2023A1515011743 to X.A.L, 2019A1515110190 to Z.X.C), Shenzhen Key Basic Research Project (JCYJ20200109115641762 to Z.X.C), Shenzhen governmental grant (ZDSYS20190902093601675 to Z.X.C), and CAS Key Laboratory of Brain Connectome and Manipulation (2019DP173024 to X.A.L and Z.X.C).

## Competing Interests

The authors declare no competing interests.

## Supplemental data

**Table. S1.**
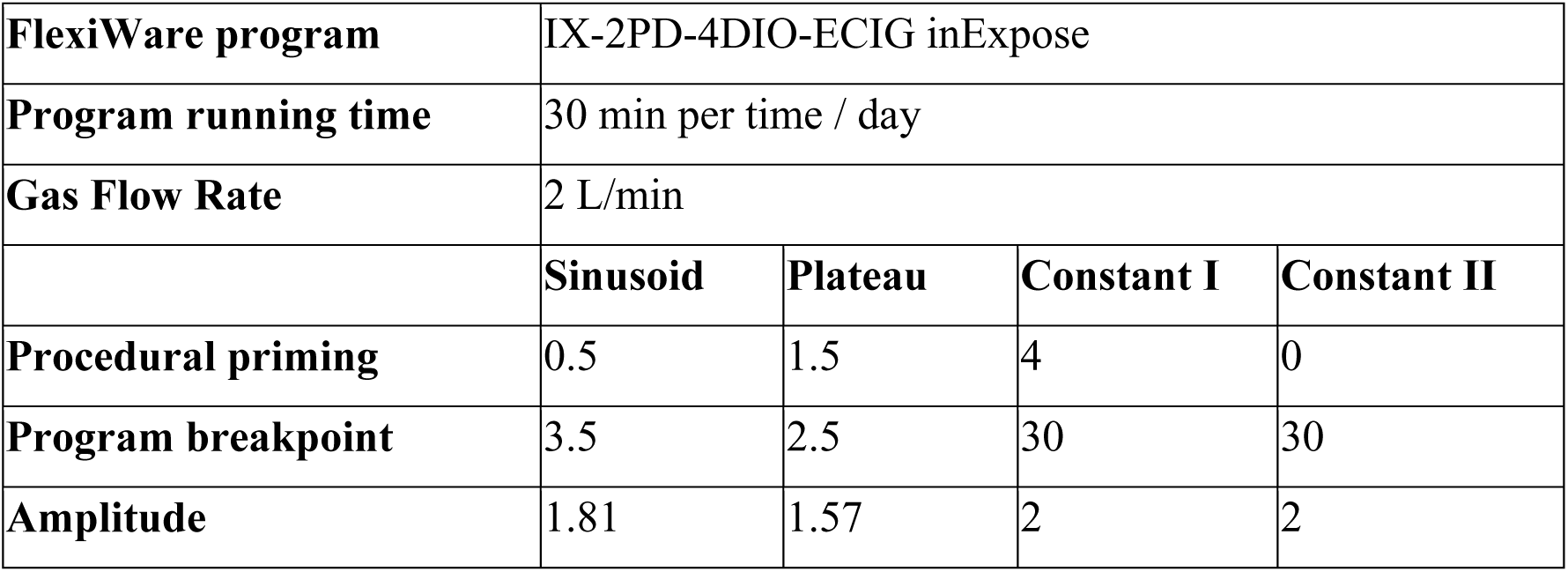
Apparatus parameters of e-cigarette vapor exposure program. The e-cigarette puff profile is configured with a combination of 3-second puff duration, 30-second puff interval, and a puff volume of 55 mL.

**Fig. S1.**
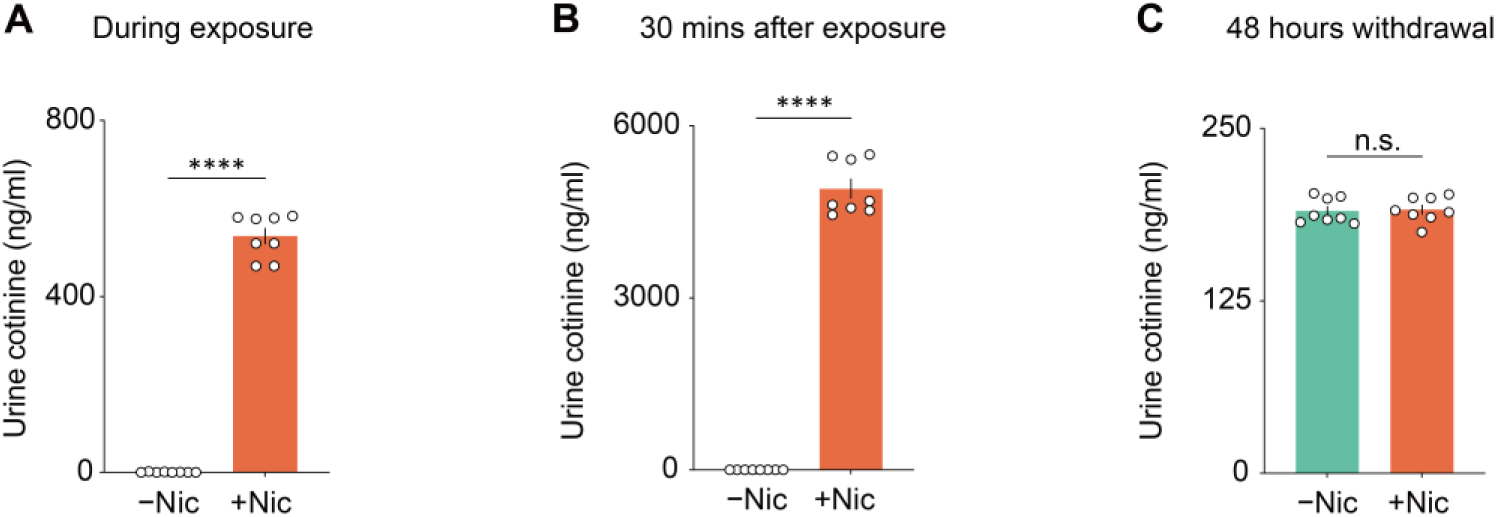
Urinary cotinine exhibits dynamic changes through measurements. **A**, The urinary cotinine concentration of the -Nic group is significantly lower than that of the +Nic group during the first day of vapor exposure. **B**, The cotinine level of the -Nic group is significantly lower than that of the +Nic group in the urine collected 30 minutes after completing vapor exposure on the first day. **C**, The result of cotinine concentration shows no significant difference between the -Nic group and the +Nic group in the urine obtained after 48 hours of withdrawal following long-term vapor exposure. (∗∗∗∗P < 0.0001.).

**Fig. S2.**
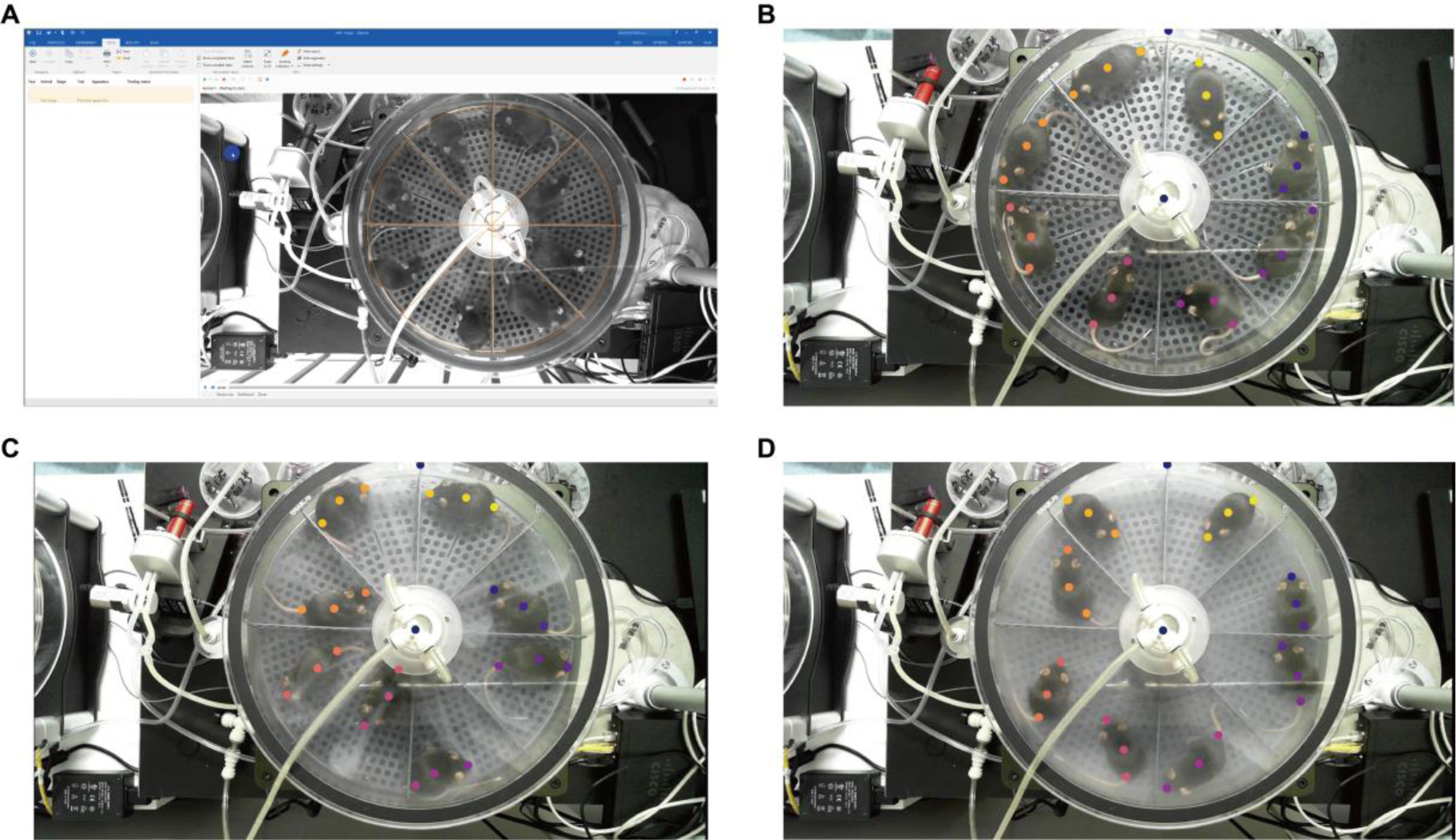
Tracking result visualization of ANY-maze and MiceVAPORDot. **A**, ANY-maze cannot detect the mice even adjusting the software parameters. **B**, MiceVAPORDot can track each mouse before smoke release. **C**, MiceVAPORDot can track each mouse during smoke release. **D**, MiceVAPORDot can track each mouse after the smoke fills the chamber.

## Notes

### Competing Interest Statement

The authors have declared no competing interest.

