## Supplemental data for "MiceVAPORDot: A novel automated approach for high-throughput behavioral characterization during E-cigarette exposure in mice"

| **FlexiWare program** | IX-2PD-4DIO-ECIG inExpose | | | |
|---|---|---|---|---|
| **Program running time** | 30 min per time / day | | | |
| **Gas Flow Rate** | 2 L/min | | | |
| | **Sinusoid** | **Plateau** | **Constant I** | **Constant II** |
| **Procedural priming** | 0.5 | 1.5 | 4 | 0 |
| **Program breakpoint** | 3.5 | 2.5 | 30 | 30 |
| **Amplitude** | 1.81 | 1.57 | 2 | 2 |

**Table. S1| Apparatus parameters of e-cigarette vapor exposure program.** The e-cigarette puff profile is configured with a combination of 3-second puff duration, 30-second puff interval, and a puff volume of 55 mL.

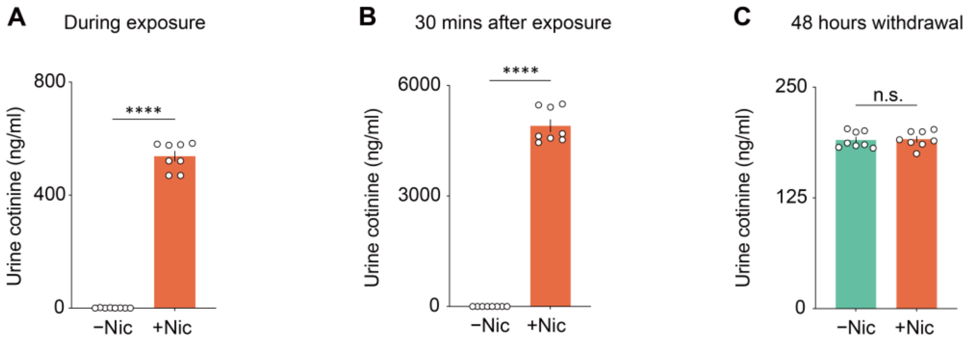


**Fig. S1| Urinary cotinine exhibits dynamic changes through measurements. A**, The urinary cotinine concentration of the -Nic group is significantly lower than that of the +Nic group during the first day of vapor exposure. **B**, The cotinine level of the -Nic group is significantly lower than that of the +Nic group in the urine collected 30 minutes after completing vapor exposure on the first day. **C**, The result of cotinine concentration shows no significant difference between the -Nic group and the +Nic group in the urine obtained after 48 hours of withdrawal following long-term vapor exposure. (*

***P < 0.0001.).

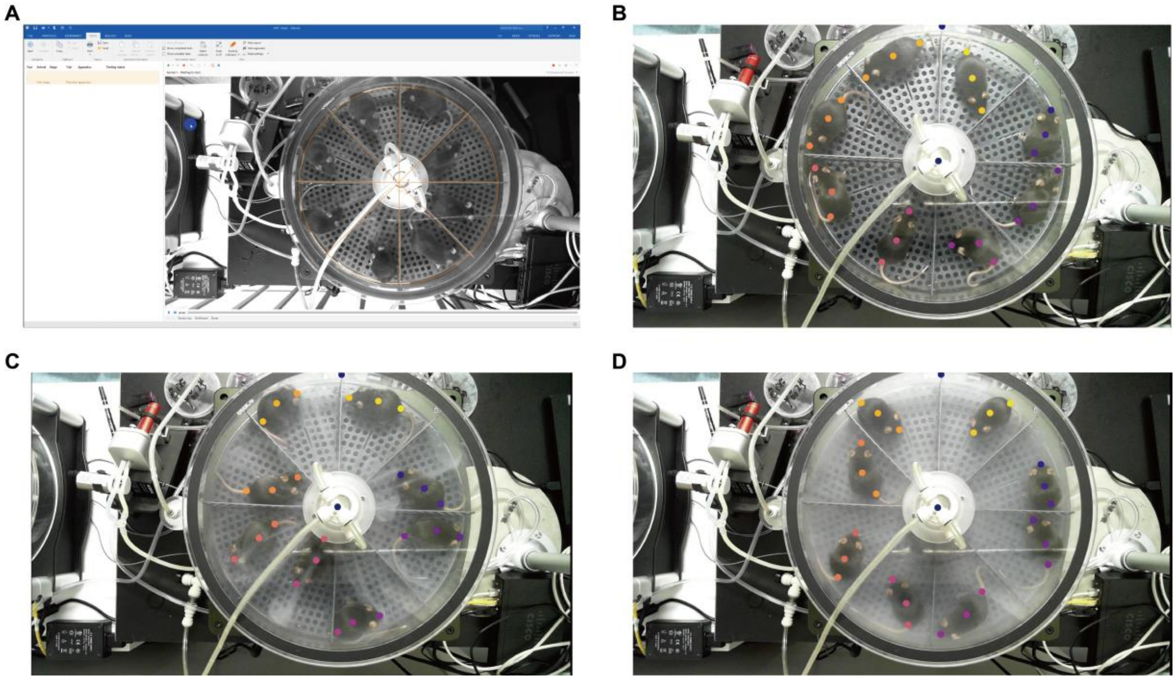


**Fig. S2| Tracking result visualization of ANY-maze and MiceVAPORDot. A**, ANY-maze cannot detect the mice even adjusting the software parameters. **B**, MiceVAPORDot can track each mouse before smoke release. **C**, MiceVAPORDot can track each mouse during smoke release. **D**, MiceVAPORDot can track each mouse after the smoke fills the chamber.
